# Identification of single nucleotide genetic polymorphism sites using machine learning methods

**DOI:** 10.1101/2023.10.19.563060

**Authors:** Mikalai M. Yatskou, Elizabeth V. Smolyakova, Victor V. Skakun, Vasily V. Grinev

**Affiliations:** Department of Systems Analysis and Computer Modelling, Belarusian State University, Minsk, Belarus; Department of Genetics, Belarusian State University, Minsk, Belarus

**Keywords:** Single Nucleotide Polymorphism, Simulation Modelling, Machine Learning

## Abstract

The paper presents an algorithm for simulation modelling of nucleotide variations in the genomic DNA molecule. To identify single nucleotide genetic polymorphisms, it is proposed to use machine learning methods trained on simulated data. A comparative analysis of the effective classical and machine learning algorithms for identifying single nucleotide polymorphisms was performed on simulated data. The most optimal method for identifying single nucleotide genetic polymorphisms in DNA molecules at various experimental noise levels is the machine learning algorithm CART.

## 1. Introduction

Genetic processes are studied using genomic sequencing experiments, which observe information on the composition of DNA and RNA molecules and their coding fragment expressions [1]. Complete genome sequencing or sequencing of only functionally significant regions of the human genome allows simultaneously identifying multiple sites of single nucleotide polymorphism (SNP), having diagnostic or prognostic significance for many human diseases [2, 3]. Statistical methods of binomial distribution, entropy-based, Fisher’s exact tests and machine learning models are used for identifying the SNPs [2, 4]. These methods are quite universal and simple for program implementation, however, are computationally expensive and difficult to be effectively applied in the analysis of experimental data with a high noise level and various experimental distortions, which are sources of gaps, repetitions, and other anomalous values [1]. Practical experimental studies use simulation modelling to select the most optimal SNP identification algorithm, test competing pipelines of analysis, and evaluate the performance of specific experimental designs for studying biophysical systems [5]. Simulation modelling is also used to generate training data for machine learning methods to directly identify SNP sites in real data from a single sequencing experiment [6]. In this case, the formation of simulated training data can have advantages in terms of accuracy and efficiency in the analysis of experimental data both with a low number of coverages and with gaps due to experimental distortions. It is expected that simulated data from a specific experiment on the human genome will provide more accurate training for machine learning SNP identification algorithms than publically available datasets.

This work presents a simulation model of the nucleotide sites in a DNA molecule and a comparative analysis of the most effective classical and machine learning SNP identification algorithms. The simulation model allows to generates datasets both for training machine learning models and for testing available SNP identification algorithms. The performance of selected SNP identification algorithms was assessed in the course of a comparative analysis on simulated sequencing data.

## 2. Simulation modelling of SNP sites

Simulation modelling of SNP sites is carried out based on experimental data, under the assumption that the main data characteristics, such as the number of nucleotide coverages, are of the beta or normal distribution [7]. Suppose a site *j* contains the reference nucleotide base *r* (nucleotides A, C, G, or T); *D* = {*b*_1_, b_2_, b_3_, b_4_} is a set of *n* reads (coverages) of nucleotide bases A, C, G or T, recorded from sequencing the site *j*; the numbers of site coverages *n, b*_1_, b_2_, b_3_, b_4_ obey the beta (Equation 1) or normal (Equation 2) distributions

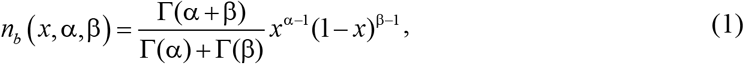

where β and α (β, α > 0) are some parameters that determine the shape of the distribution curve; Γ is the gamma function;

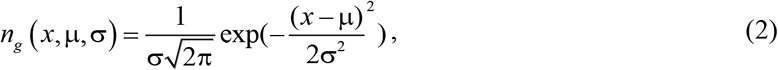

where μ and σ are parameters of mathematical mean and standard deviation.

The idea of modelling is to randomly generate *N*_SNP_ positions of SNP sites in the sequence of the considered molecule *S*, consisting of *N* nucleotide sites, for each of which the numbers of coverages *n, b*_1_, *b*_2_, *b*_3_, *b*_4_ are reproduced according to the beta or normal distributions in the defined range [*n*_min_; *n*_max_]. For a non-reference site *j*, the total number of coverages *n* is modeled, then the numbers of coverages for the reference *b*_*Ref*_ and non-reference *b*_*nRef*_ nucleotides are generated from the resulting *n*. Nucleotide coverages for the SNP site are modeled similarly. It is assumed that there are coverages of no more than two different nucleotide bases on the site. For a comprehensive study of SNP identification algorithms, the addition of Gaussian noise with parameters μ = 0 and σ_*l*_ = *q*_l_ · *b*_*l*_, *l* = 1-4 (indexed nucleotides A, C, G, and T), to the numbers of nucleotide coverages *D* were implemented (Equation 3)

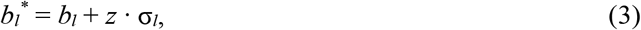

where *z* is the realization of a standardized normal random variable, *q*_*l*_ > 0. Varying the parameter *q*_*l*_ changes the level of experimental noise, namely, it regulates the informativeness of the useful signal, which allows comprehensively studying the effectiveness of selected SNP identification algorithms and recreate special experimental conditions.

The proposed simulation algorithm reproduces datasets as close as possible to experimental conditions, given by the numbers of site coverages and the laws of their distributions, the number of SNP sites. The flow diagram of the algorithm for modelling SNP sites is shown in figure.

**Figure. 1.**
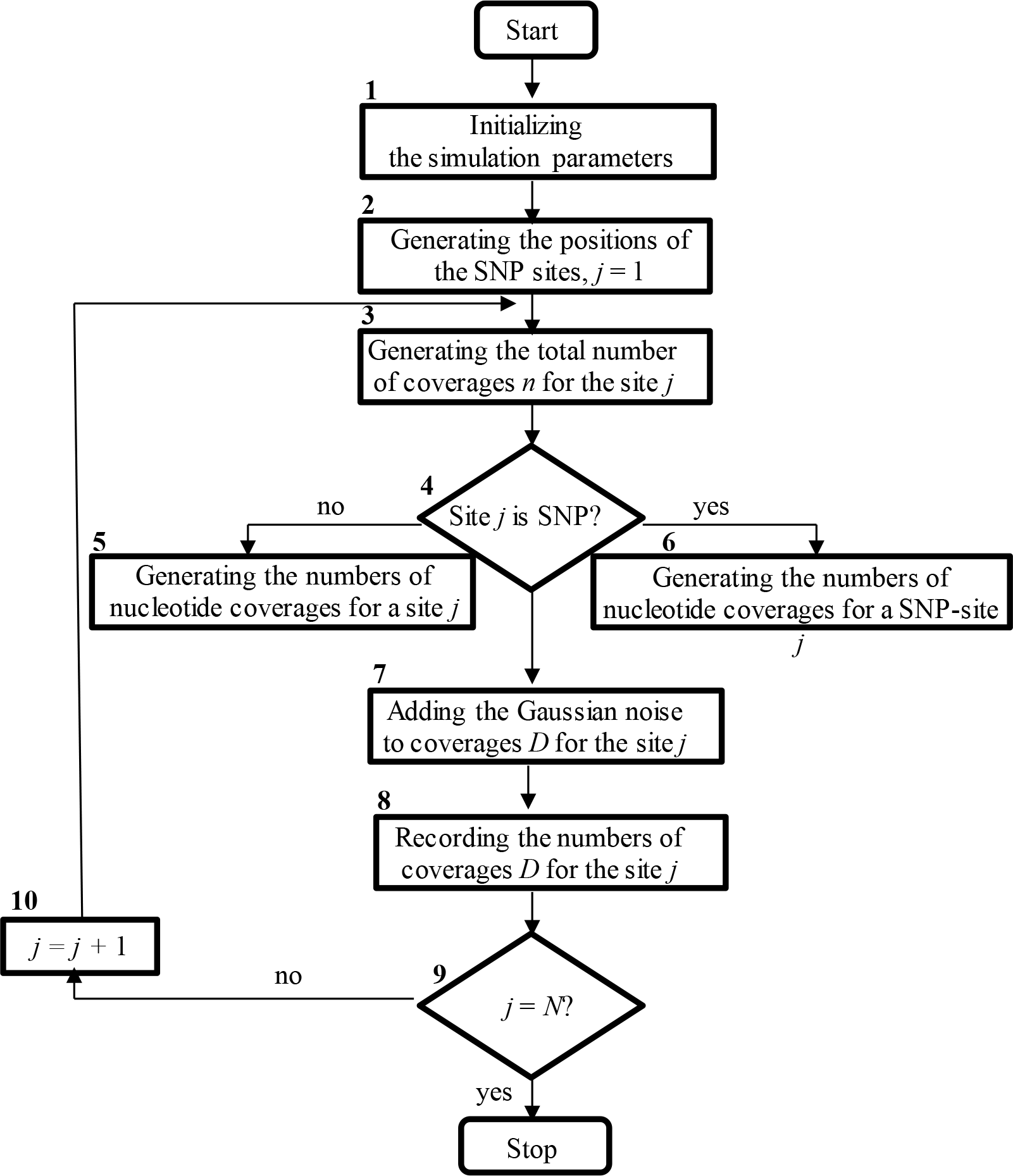
Flow diagram of the algorithm for simulation modelling SNP sites.

### Algorithm.

Step 1. Initialize the model parameters *N, N*_*SNP*_, *n*_min_ and *n*_max_, α and β (or μ and σ) (Figure, block 1). Parameters α and β (or μ and σ) are given for distributions of the numbers of site coverages *n, b*_1_, *b*_2_, *b*_3_, *b*_4_.

Step 2. Generate the SNP site positions *L* = {*l*_1_, *l*_2_, …, *l*_NSNP_} in the sequence *S* according to the uniform discrete distribution in the interval [1; *N*] (block 2). Set the position index *j* = 1.

Step 3. Gamble the total number of reads *n* on the current site *j* as a realization of a random variable of the beta or normal distribution with experimentally extracted parameters (block 3).

Step 4. Check if the site *j* is SNP. Accordingly go to step 5 or 6 (block 4).

Step 5. Generate the numbers of nucleotide coverages *b*_1_, *b*_2_, *b*_3_, *b*_4_ by the beta distribution with experimentally assessed parameters for non-SNP sites (block 5). Go to step 7.

Step 6. Generate the numbers of nucleotide coverages *b*_1_, *b*_2_, *b*_3_, *b*_4_ by the beta distribution with experimentally assessed parameters for SNP sites (block 6).

Step 7. Add the Gaussian noise to the number of nucleotide coverages *b*_1_, *b*_2_, *b*_3_, *b*_4_ for a site *j* (Equation 3, block 7).

Step 8. Record the simulated characteristics of the site *j* to a data file (block 8).

Step 9. Check the termination condition of the simulation algorithm (block 9). If all sites in the sequence are simulated, i.e. *j* = *N*, then stop the simulation. Otherwise *j* = *j* + 1 (block 10) and go to step 3.

## 3. Machine learning algorithms

To apply machine learning algorithms, it is necessary to form a set of features charactering a nucleotide site. It was decided to use 4 features: *X*_1_ – the number of coverages of the reference nucleotide, *X*_2_-*X*_4_ – the numbers of coverages for non-reference nucleotides sorted in descending order. The data are normalized to the total number of site coverages *n*. Taking into account the limited number of 4 features, and the binary classification problem (SNP and non-SNP site classes) to be solved, it is preferable to test basic machine learning methods, such as Conditional Inference Trees (CIT), Classification And Regression Tree (CART), Support Vector Machines with a linear separating function (SVM), and Extreme Gradient Boosting (XGBoost). Let’s take a closer look at the selected methods.

CIT. The algorithm is based on the use of the Strasser and Weber statistical test [8]. Binary partitioning at a tree node is carried out according to a feature *X*_*j*_, for which the main and alternative hypotheses about the statistical relationship with the output variable *Y* are formulated. To test the hypothesis, the Strasser and Weber permutation test is used and *p*-values are calculated. The feature for which the *p*-value is minimal is selected as a partition node *X*_*j*_. The advantage of the algorithm is the use of a statistical criterion and relatively high accuracy among classical machine learning algorithms.

CART. Binary splitting in a node of a tree is carried out according to a feature *X*_*j*_, the criterion for splitting a node is the Gini index, the threshold for splitting the feature is selected based on the minimum of the Gini index [9]. The advantages of the algorithm are versatility and compactness.

SVM. The method is designed to find optimal, in a certain sense, data classification functions (decision functions) [10]. The advantage is simplicity and efficiency in separating two-class problems. XGBoost. The method is based on a gradient boosting algorithm on regression decision trees that approximate the negative gradient functions constructed from the samples of the training dataset, the result of which determines the contributions of *m* weak classifiers to the overall classifier [11]. The sample drops to the class whose probability is maximal. The advantage of the algorithm is its high accuracy and speed of calculations (compared to other ensemble algorithms).

## 4. Organization of a computational experiment

In our computational experiment the machine learning models were trained on specially simulated datasets and then the comparative analysis of the classical and machine learning SNP identification algorithms was performed on other generated datasets with varying levels of the added Gaussian noise.

The machine learning models of CIT (the R function *ctree* of the package *party*), CART (the R function *rpart* of the package *rpart*), SVM (the R function *svm* of the package *e1071*) and XGBoost (the R function *xgboost* of the package *xgboost*) were trained on synthetic data simulated using the beta distribution with no adding any Gaussian noise. A training dataset contained 40,000 sites, of which 20,000 were SNPs.

We included in the comparative analysis two most effective existing SNP identification algorithms – the binomial distribution and entropy-based tests [2, 4]. An efficient software implementation of the binomial distribution test (BDT) has been developed, a feature of which is the automation of the selection of a threshold value when identifying SNP sites. It is proposed to use the value 10^-*k*^ as a threshold value of probabilities, where *k* is the average number of site coverages estimated from the simulated or experimental dataset. The published software implementation is used as an entropy-based test (EBT) [4]. Thresholds in identifying SNP sites are the entropy *E* > 0,21 and the *p*-value < 0,5.

For a comprehensive study of SNP identification algorithms datasets were simulated taking into account the addition of varying Gaussian noise. Two groups of datasets were generated: 1) the parameter values for the reference and non-reference channels of the site *q*_*R*_ and *q*_*nR*_ were assumed equal and varied from 0 to 0,6 – these datasets allow to investigate the influence of the increasing noise level in the nucleotide channels of coverages on the accuracy of the SNP identification algorithms; 2) the parameters *q*_*R*_ = 0 and *q*_*nR*_ varied from 0,5 to 2,0 – these datasets are for investigating the influence of increasing noise level in the non-reference channel on the accuracy of the algorithms. Datasets of 20,000 sites were simulated, each with 20 randomly generated SNP sites. The number of datasets for each parameter combination was 3.

The performance of the SNP identification algorithms was evaluated using the standard classification measures for unbalanced classes, such as *Precision, Recall* and score *F*_1_, characterizing the properties of the algorithms accept false positive (non-SNPs as SNPs, *Precision*) and false negative (SNPs as non-SNPs, *Recall*) events, and their combined contribution the score *F*_1_ [12].

In the course of the work, R-functions were developed that implement various stages of simulation modelling and SNP identification algorithms. It is proposed to integrate the developed functions into a dedicated R-package that can be used to model synthetic datasets, according to a concrete experiment, in order to comprehensively test and select the best algorithms for identifying SNP sites, as well as for generative data modelling to train identification algorithms based on machine learning models.

## 5. Results

Based on the sets of simulated data, we conducted a comparative analysis of the most effective SNP identification and machine learning algorithms, trained on simulated data. The results of the comparative analysis of the algorithms are collected in table.

**Table 1.** Accuracy of SNP identification algorithms based on the score *F*_1_.

| $q_R; q_{nR}$ | $F_1, \%$ | | | | | |
| --- | --- | --- | --- | --- | --- | --- |
|  | BDT | EBT | CIT | CART | SVM | XGBoost |
| 0; 0 | 91,9 (0,8) | 97,6(1,4) | **100 (0)** | **100 (0)** | **100 (0)** | **100 (0)** |
| 0,2; 0,2 | 91,8 (2,1) | 96,0 (0,8) | **99,0 (0,8)** | 98,3 (0,9) | 98,3 (0,9) | 98,3 (0,9) |
| 0,4; 0,4 | 84,1 (1,8) | 82,3 (3,2) | 47,4 (0,5) | **88,8 (3,1)** | 2,6 (3,0) | 44,9 (1,5) |
| 0,6; 0,6 | **82,7 (4,5)** | 79,3 (2,3) | 19,0 (1,1) | 81,1 (1,7) | 1,6 (1,6) | 17,6 (1,3) |
| 0,0; 0,5 | 87,0 (3,0) | 92,2 (1,9) | **92,6 (2,4)** | 91,8 (1,7) | 92,0 (2,8) | 91,8 (1,7) |
| 0,0; 1,5 | 75,8 (2,9) | 77,1 (3,4) | 85,6 (2,8) | **88,2 (4,0)** | **88,2 (4,0)** | **88,2 (4,0)** |
| 0,0; 1,5 | 72,6 (2,5) | 74,7 (2,1) | 87,0 (3,5) | 92,1 (2,7) | **92,3 (1,6)** | 92,1 (2,7) |
| 0,0; 2,0 | 56,7 (0,6) | 59,5 (1,6) | 72,9 (1,7) | **88,9 (2,0)** | 85,6 (2,8) | **88,9 (2,0)** |

*Note.* The standard error of the mean is indicated in parentheses.

Datasets 1. On the dataset with no adding Gaussian noise, the highest accuracy of the score *F*_1_ (100 %) is obtained for the machine learning methods. The accuracy of EBT (97,6 %) is higher than that of BDT (91,9 %). Wen increasing noise in the data from *q*_*l*_ = 0,2 to 0,6, the accuracy of the BDT, EBT and CART algorithms decreases to 80-82%, and for the machine learning models CIT, SVM and XGBoost – to 18 % and lower. The poorest accuracy, when noise increases from 0.4 and higher, is observed for the SVM model (1,6-2,6 %).

Datasets 2. When the noise in the non-reference channel increases from *q*_*nR*_ = 0,5 to 2.0, the accuracy of classical algorithms decreases significantly to 57-60%, and of machine learning algorithms to 73-89%. The CIT model has the lowest accuracy among machine learning methods when noise increases from *q*_*nR*_ = 1,5 (73%).

These results allow to conclude that for non-noisy data it is preferable to use machine learning algorithms. When data are uniformly noisy in the nucleotide channels, it is advisable to use classical algorithms and the CART model; when non-reference channels are noisy then the machine learning algorithms should be applied. The poor classification accuracy for the CIT algorithm at higher noise levels can be explained by the deterioration of the statistical properties of the samples under consideration, which is critical for statistical algorithms.

## 6. Conclusions

It is proposed to use machine learning methods trained on simulated data to identify the single nucleotide genetic polymorphism sites. An algorithm has been developed for simulation modelling of single nucleotide sites in the genomic DNA molecule, based on the generation of random events according to the beta or normal distribution. A comparative analysis of the most effective classical and machine learning algorithms for identifying single nucleotide polymorphism sites, trained on simulated data, was performed. Using examples of non-noisy data – the best methods are of machine learning; with increasing the noise level – the binomial distribution and entropy-based tests and CART. When adding noise to non-reference channels, the best methods are of machine learning – CART, SVM and XGBoost. The conducted research allows to conclude that the most optimal method for identifying single nucleotide genetic polymorphisms at various experimental noise levels is the machine learning algorithm CART.

## Notes

### Competing Interest Statement

The authors have declared no competing interest.

